# Ingestive behaviour of grazing ruminants: meta-analysis of the components linking bite mass to daily intake

**DOI:** 10.1101/705665

**Authors:** M. Boval, D. Sauvant

**Author notes:** Corresponding author: Maryline Boval.

## Abstract

This meta-analysis shed light on the quantitative adaptive responses of feeding behaviour of Cattle (C) and Small Ruminants (SR), facing variations of sward characteristics, notably of sward height (SH, 18. 7 ± 13.9 cm) and herbage bulk density (HBD, 1.73 ± 1.30 kg DM / m^3^). All responses expressed a plateau stressing an adaptive limit with extreme values of SH and HBD. The minimum plateau of BR (46.9 ± 14.6 min-1) is around 40 min-1, while IR values (different for C and SR, respectively 69.1 ± 38.1 vs. 99.9 ± 45.7 g/min/kg BW) ranged between a minimum and maximum plateau around 50 and 100 g/min/kg BW. Two other pasture management factors affect IR, namely forage allowance (10.16 ± 6.0, DM % BW) and daily proportion of time spent grazing (0.30 ± 0.08). The results obtained confirm the specifically key role of BM (1.80 ± 127 mg DM/kg BW) on IR. The regressions are IR=145 (1-exp (-b BM), b being equal respectively for C and SR and C to 0.44 vs. 0.54. This literature review has also revealed fundamental differences in behaviour between C and SR although no study to date has attempted to compare them simultaneously. SR have to chew more (2.7 ± 1.2 vs. 1.6 ± 0.5 JM/bite) to ingest the same amount of DM per bite than C, expressed in relation to BW, which allow them to ingest slightly quickly.

**Implications:** This article, following the previous one of Boval and Sauvant (2019), proposes a quantitative appraisal of the ingestive behaviour of grazing ruminants, based on studies published over 40 years, as well as well robust average values and relationships, considering inter- and intra-study effects and animal species specificities. This knowledge should contribute to a better overall understanding of the behavioural adaptation of ruminants at pasture, to the identification of key threshold values and appropriate parameters of interest to be considered, and to improve the efficiency and sensitivity of automatic devices, which are booming in the context of precision livestock farming at pasture.

## Introduction

Knowledge of ingestive behaviour (IB) is determinant to better understand the strategies of animals for feeding in order to improve their management, whatever the feeding context. Ingestive behaviour determines the nutrient supply to ruminants and thus has a significant impact on performance and feed efficiency, which are essential for increasing the profitability of livestock (Llonch *et al*., 2018; Shalloo *et al*., 2018). Moreover, a ruminant’s robustness partly comes from its ability to adapt IB to the diversity of resources to be grazed. In addition, chewing behaviour provides information about digestive comfort and indicators of appetite, gut health and welfare. Numerous studies have focused on the IB of grazing ruminants. However, beyond the problems linked with the diversity of the methodologies applied, the items measured are very heterogeneous across publications. Likely for this reason, no synthetic statistical interpretation of published IB data for ruminant grazing has been carried out and published so far. However, there is a need to better understand the various aspects of animal IB, as it is now becoming possible to measure some of them in field conditions thanks to advances in electronic/computer technologies (Anderson *et al*., 2014; Fogarty *et al*., 2018). Indeed, animal behaviour documented by tools employed in precision livestock farming, such as sensors, video cameras, accelerometers or pedometers, should greatly help in designing efficient management strategies for livestock production systems (di Virgilio *et al*., 2018). A recent meta-analysis focused on bite mass (BM) and its main determining factors linked to major animal and sward characteristics (Boval and Sauvant, 2019). In this current paper, we will analyse the components affecting BM which are determining for intake rate and daily dry matter intake. Some studies differentiate between jaw movements due to harvesting forage and those processing the forage before swallowing. We have therefore attempted to better understand the function of these different types of jaw movements.

## Material and methods

### Literature review and dataset construction

This meta-analysis was carried out by considering published studies measuring components of the feeding behaviour of ruminants (cattle, sheep or goats) at pasture in various production systems (milk or meat) and various climatic contexts. The search for the literature was carried out using Web of Science, Science Direct, EDP Sciences and Cambridge Journals and using the reference lists cited by some reviews on the subject.

For each publication, we have integrated experiments and treatments for which there were documented values of at least one of the following criteria: BM, biting rate (BR), intake rate (IR), grazing and ruminating times (GT and RT, respectively, min/day) and data related to dry matter intake (DMI) and BW gain. In some publications, total jaw movements and chews were also measured, and therefore were included.

### Intermediary calculations

For all the characteristics, we have harmonized the units within the whole dataset. Afterwards some components were also expressed per kg of BW, such as IR, in order to analyse the whole dataset including the maximum degrees of freedom (with data coming from different species and types of domestic ruminants). Considering the BM calculation, Boval and Sauvant (2019) have shown that BM can be divided by the BW^1^. When it was possible, the number of chews was calculated, considering that jaw movements (JM) = chews + bites (Galli *et al*., 2017; Mulvenna *et al*., 2018). When JM are expressed per bite, the number of JM/bite cannot be lower than 1 (i.e. one bite). For each publication retained in the database, the following information was recorded: the animal characteristics (breed, sex, age) as well as the forage characteristics (species, herbage mass, surface sward height and herbage bulk density, morphological and chemical composition, etc.). Information related to the experimental conditions (at pasture or in other environments) and to the methods used to measure feeding behaviour and forage characteristics was also recorded. The season (dry or rainy), latitude and longitude have been précised as well for each experiment intra-publication, by using Köppen–Geiger classification (Peel *et al*., 2007).

### Treatment encoding

Beyond specific codes assigned to each publication and to each experiment, additional codes were applied to identify specifically the factors of variation tested in the papers: the forage species, sward height, herbage bulk density or herbage allowance and the animal species. All of these codes were specific to the factors of variation studied in the publication; therefore, not all rows have values in the corresponding columns. For some experiments, in addition to the intra-experimental factors, some key criteria varied significantly, although they were not the factors tested intra-experiment. In this case, another code was added to specify these criteria, as a secondary factor of variation. For example, we identified experiments for which the intra-experiment sward height varied largely despite not being announced as a factor in the publications, but which can then be considered as a factor of variation for 62% of papers instead of the 32% we had identified at first approach.

The final database included 98 publications (npub), 269 experiments (nexp) and 905 treatments (n). The list of the references used to build the database is presented in the Annex.

### Statistical analysis

Statistical analysis of the data was performed by meta-analysis according to the recommendations of Sauvant *et al*. (2008). In particular, inter- and intra-experiment variations were split to study in the intra-experiment relationships between variables considered two by two, and successively through the various factors of variations. The numbers of data different from one variable to another explain why the interpretation must be achieved considering the variables 2 by 2.

## Results

### Statistical parameters of the ingestive behaviour components

The statistics of the components of feeding behaviour were calculated for cattle and small ruminants (Table 1) and according to BW when it allowed pooling of data for both species. Among the data collected in the papers analysed, BR was one of the most documented components, as well as its reverse, namely the time spent per bite. These two components have a log-normal distribution and do not differ significantly between species.

**Table 1.** Number and mean values, standard deviation, minimums and maximums of the feeding behaviour components collected in the publications

| | Mean ± SD | Min–max | n | Normality<sup>1</sup> |
| --- | --- | --- | --- | --- |
| **BM (mg DM/BW)** | 1.80 ± 1.27 | 0.107–7.41 | 581 | N(L) |
| Small ruminants (g DM/B) | 0.11 ± 0.09 | 0.09–0.63 | 117 | N(L) |
| Cattle (g DM/B) | 0.77 ± 0.65 | 0.05–4.00 | 458 | N(L) |
| **BR (bites/min)** | 46.9 ± 14.6 | 11.2–106.7 | 560 | N |
| **JM/bite** |  |  |  |  |
| Small ruminants | 2.73 ± 1.22 | 1.0–6.5 | 45 | N |
| Cattle | 1.60 ± 0.54 | 1.13–5.04 | 97 | N |
| **Chews/bite** |  |  |  |  |
| Small ruminants | 1.79 ± 1.44 | 0.20–6.20 | 35 | N |
| Cattle | 0.59 ± 0.52 | 0.12–3.36 | 92 | N |
| **IR (g DM/min)** |  |  |  |  |
| Small ruminants | 4.34 ± 2.0 | 0.5–11.3 | 91 | |
| Cattle | 27.6 ± 18.9 | 1.3–146.3 | 339 | |
| **IR (mg/min/kg BW)** |  | 2.70–274.1 | 415 | N(L) |
| Small ruminants | 99.94 ± 45.7 | 12.0–274.1 | 64 | |
| Cattle | 69.1 ± 38.1 | 2.70–248.8 | 349 | |
| **Bite/g DMI** |  |  |  |  |
| Small ruminants | 15.8 ± 17.7 | 1.6–105.0 | 85 | N(L) |
| Cattle | 2.5 ± 2.1 | 0.3–21.0 | 316 | N(L) |
| **GT (min)** | 519 ± 146 | 138–1080 | 293 | |
| **RT (min)** | 366 ± 129 | 31–574 | 140 | |
| **IT (min)** | 459 ± 190 | 86–955 | 149 | |
| **DMI (%BW)** | 2.96 ± 1.10 | 0.45–8.0 | 248 | |

BM = bite mass; B = bite; BR = bite rate; JM = jaw movements; IR = intake rate; GT = grazing time; RT = ruminating time; IT = idling time; DMI = dry matter intake.
<sup>1</sup> Normality of the distributions: N = non-normal; L suggests a log-normal asymmetric distribution.

Jaw movements associated with bites were registered in only about 20% of cases, but with sufficient data for each species.

Intake rate was less much documented in the papers than BR, and distribution of the data is quite close to a Gaussian law. Otherwise, IR is largely different between cattle and small ruminants, contrary to BR (Table 1). Even when IR was expressed per kg of BW, the difference between species remained but the value in this case was a little higher for small ruminants (Table 1).

### Modelling factors of variation of the ingestive behaviour components Bite rate

The influence of sward height (SH) on BR was evaluated from experiments that studied SH impacts. There is a negative curvilinear relation (Figure 1a) between BR (bites/min) and SH (18.7 ± 13.9 cm)). Under a threshold of SH of 15–20 cm, there appears to be an acceleration of BR. The intra-experimental regression is:

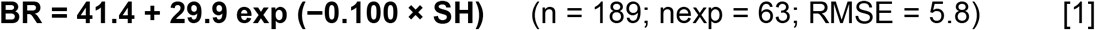

**Figure 1.**
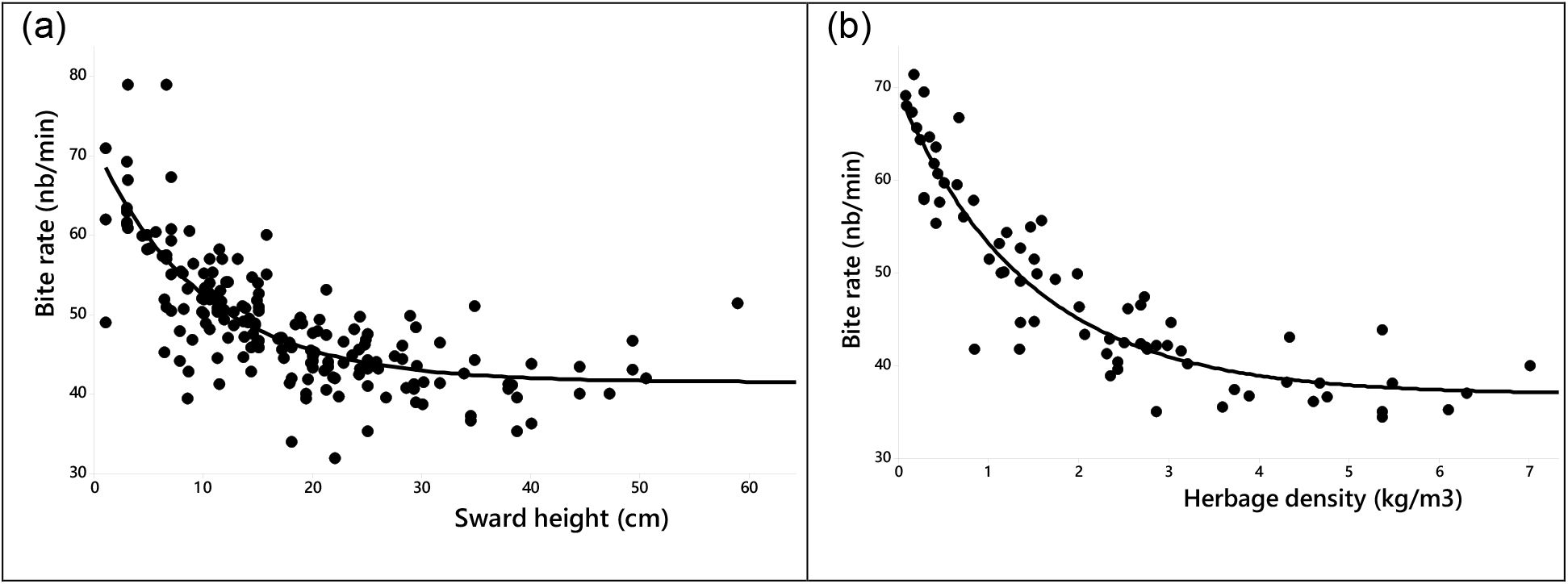
Intra-experiment relationships between bite rate and sward height (a) or herbage bulk density (b).

The impact of apparent forage density (HBD, 1.73 ± 1.30 kg DM/m^3,^) on BR was assessed from experiments that tested the impacts of HBD variations. There is a negative exponential intra-experiment relationship between BR (bites/min) and HBD (Figure 1b); the regression is:

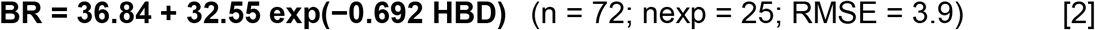

An acceleration of BR occurs when the HBD decreases below a threshold between 2 and 3 kg DM/m^3^. The regression in Equation 2 is a little more accurate than Equation 1 (RMSE = 3.9 vs 5.8).

### Intake rate

#### Impact of SH

When considering the experiments dealing with SH variations, there is a positive curvilinear intra-experiment effect of SH on IR (mg DM/kg BW/min), with an asymptotic value around 100 mg DMI/min/kg BW and a rapid decline in IR under a threshold SH of about 15–20 cm (Figure 2a). The intra-experiment relationship between both variables is:

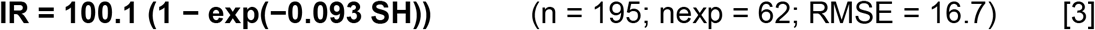

**Figure 2.**
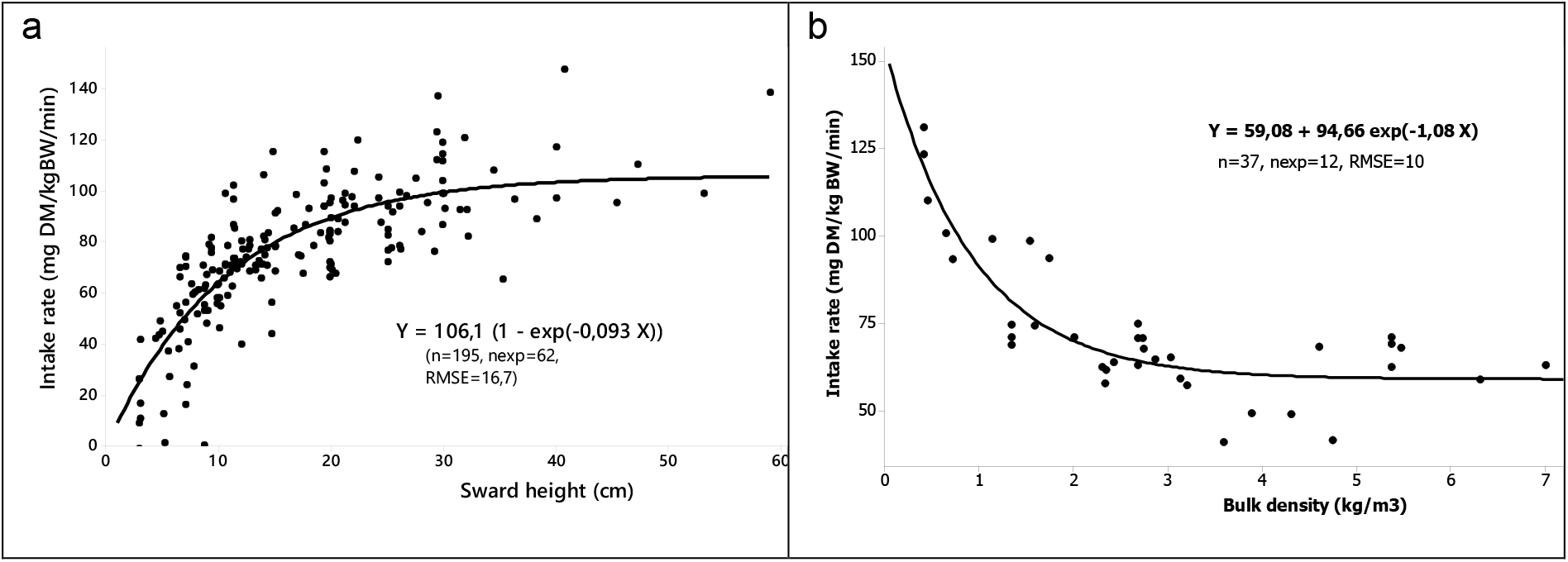
Impacts of sward height (cm) (a) and herbage bulk density on intake rate (mg DM/kg BW/min) (b).

It appears that for three publications (Black and Kenney, 1984; Mezzalira *et al*., 2014 and 2017) and nine experiments, the response of IR to SH is clearly curvilinear, exhibiting a maximum value of IR followed by a decreasing IR with increasing SH (Figure S1). In these papers, the maximum values of IR ranged between about 115 and 160 mg DM/kg BW while the corresponding values of SH ranged between about 10 and 30 cm.

#### Impact of HBD

As seen for BR (Figure 2b), there is an increase of IR when HBD is lower than a threshold of 2–3 kg DM/m^3^. The intra-experiment regression is:

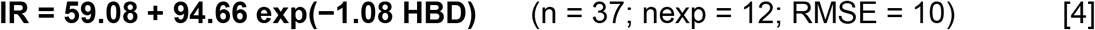

#### Interaction between SH and HBD

As mentioned, Equations 3 and 4 were calculated on datasets issued from experiments that considered variations in SH and HBD, respectively, as experimental factors. As the number of data with SH and HBD is fairly high, another approach was performed to study, within publications, the effect of interactions between SH and HBD on IR (mg DM/kg BW/min). An intra-publication significant quadratic regression was calculated:

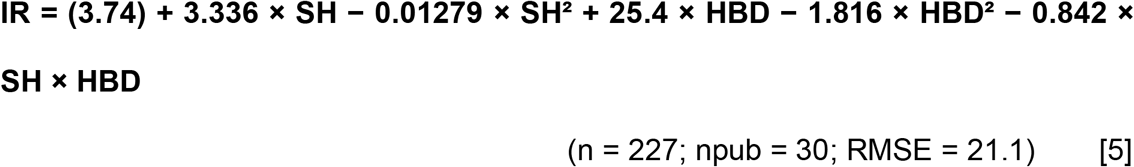

The three quadratic terms of this regression are highly significant, stressing the interaction between SH and HBD. Figure 3 shows the trace of this regression and illustrates the interaction with SH on the X-axis and HBD corresponding to the successive lines of iso-HBD. The thickness of the lines is proportional to the frequency of the observed situations. The interaction appears concretely in Figure 3: when SH < about 20 cm, its negative influence on IR is compensated by an increase of HBD. Beyond the threshold value of HBD around 2–3 kg DM/m^3^, the influence of SH almost disappears. Otherwise in Figure 3, it can be seen that HBD has no effect on IR when SH is around 1–20 cm. Over this threshold of SH around 15–20 cm, the influence of HBD tends to be negative on IR which is then impacted mainly by the variations of SH.

**Figure 3.**
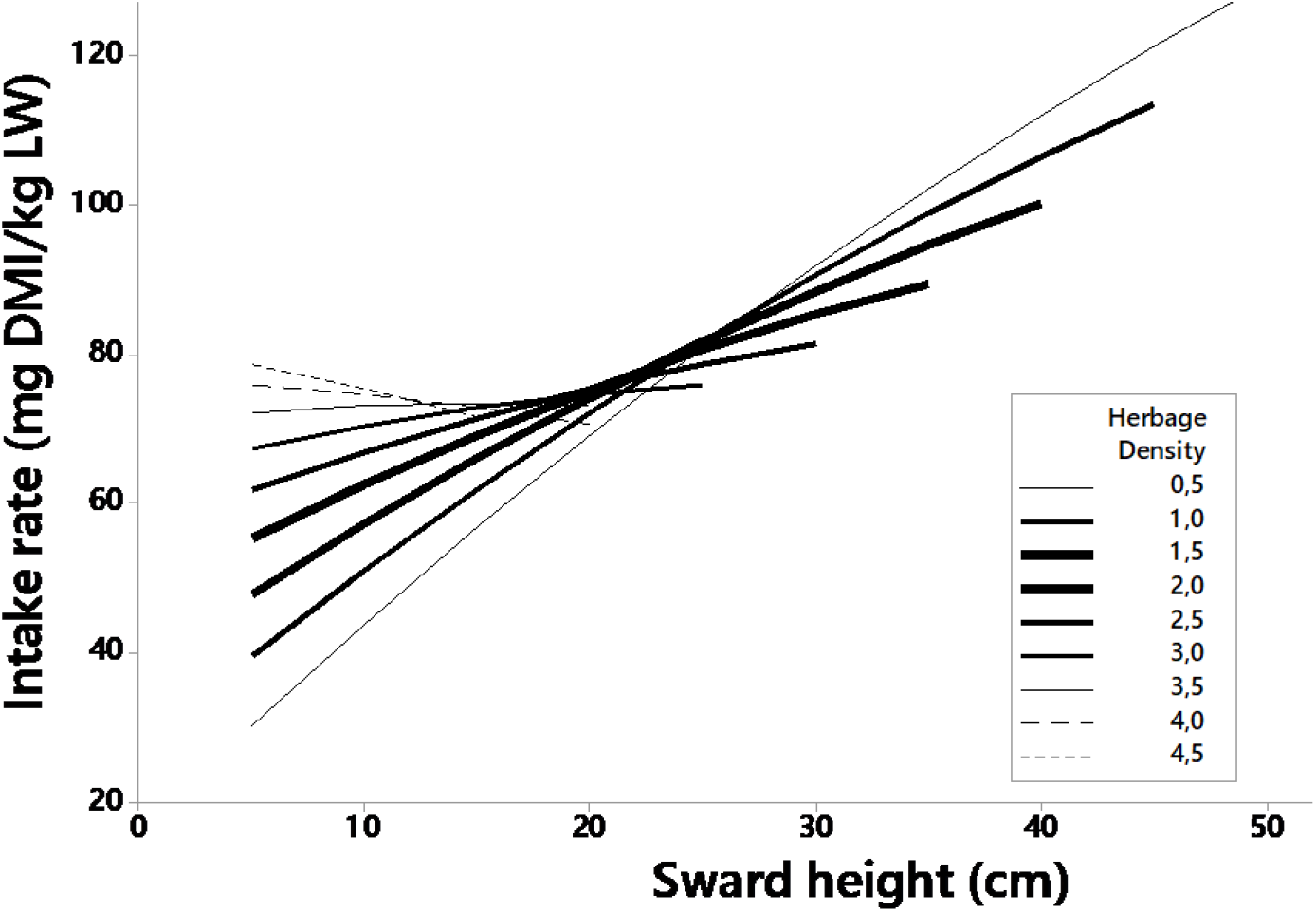
Interactions between sward height (cm) and herbage bulk density (kg DM/m^3^) on intake rate (mg DM/kg BW/min).

#### Influence of stem and leaf mass

The stem mass (SM = 1.41 ± 0.80), when leaves are available, determines IR (Figure S2) according the following intra-experiment regression:

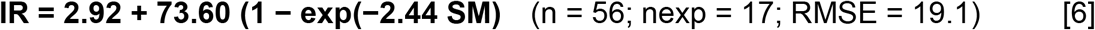

The leaf mass (LM; 1.17 ± 0.74 t DM/ha) also affects IR according to the following equation:

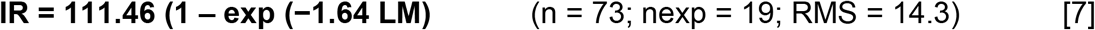

Figure S2 presents the two regressions and illustrates the fact that LM explains a large range of IR, from 0 to 111 mg DM/kg BW/min, while for SM, the equivalent range is only 48 mg DM/kg BW/min. For the lower values of SM, there is a great variability of IR, which is only due to the impact of leaf growth. Thus, it was decided to remove these low values of SM. Figure S2 shows also that the plateau is achieved for SM beyond the threshold of about 1 t DM/ha, illustrating that the continued growth of stems does not affect IR. In contrast, leaf growth goes on impacting IR, without any precise threshold of LM/ha.

### Impacts of grazing management factors

The effect of herbage allowance (HA) on IR was analysed for experiments excluding continuous grazing. It appears that IR decreased when HA increased, until a minimum plateau close to 40 mg/kg BW/min (Figure 4a). When HA decreased under a value of around 10% BW, IR increased rapidly until values close to 100 g/kg BW/min. The values under 10% of BW come partly from experiments where the access time was only 1 h or even less (Figure 4a). This response of IR is mainly due to the increase of BM.

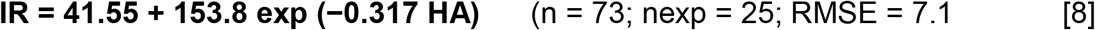

**Figure 4.**
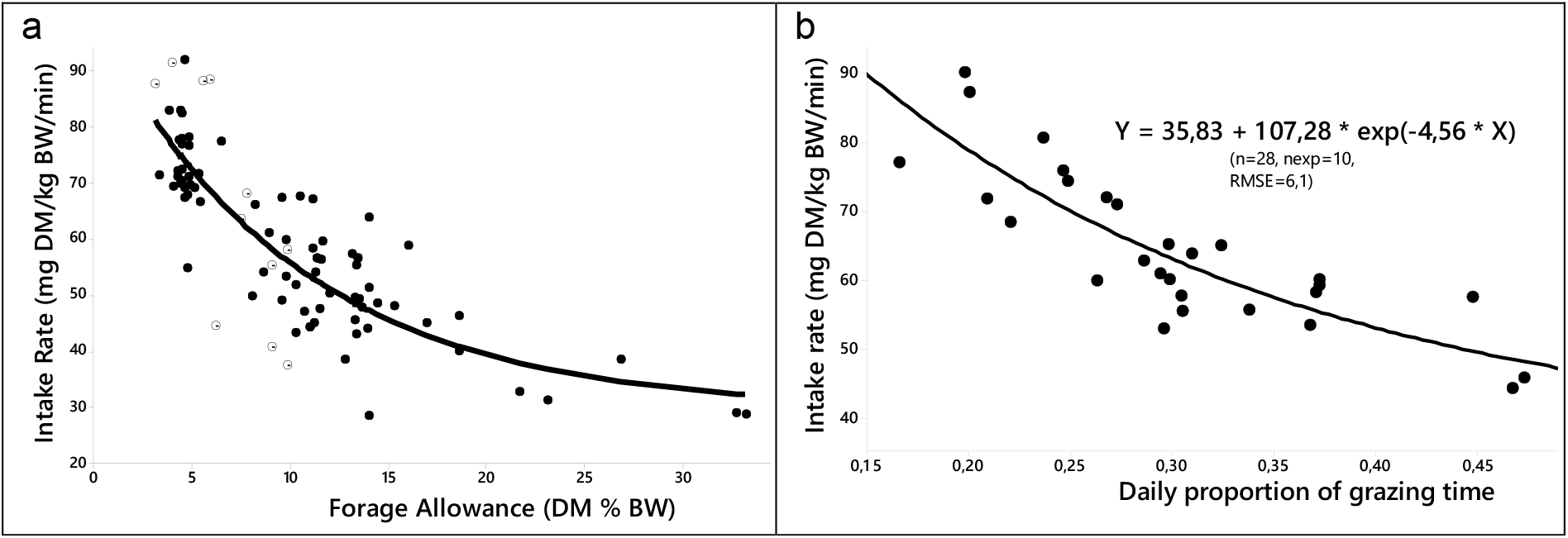
Effect of forage allowance (a) and proportion of time spent grazing (b) on intake rate (mg DM/kg BW/min).

Another major factor of grazing management is the access time. In the database, only 10 experiments were focused on this aspect. The response of IR is negatively related to the daily proportion of time spend grazing (pGT, 0.30 ± 0.08, Figure 4b):

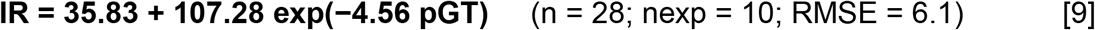

For the same dataset, there was a trend of reducing the level of DMI/BW when pGT decreased: −1.36 ± 0.81 g DMI/kg BW (*P* = 0.11) per 0.1 decrease of pGT. For a part of this dataset (three experiments and nine treatments), the BM was measured and it increased significantly when pGT decreased (−7.1 ± 1.3 g DM/kg BW per unit of pGT). In contrast, BR was not influenced by pGT in this dataset.

### Grazing time

When GT is not limited, there is a negative relationship between SH and GT (Figure S3), and the intra-experiment regression between the two parameters is:

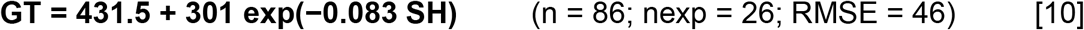

The number of data is not sufficient to study the influence of HBD on GT, or on rumination and idling times.

### Interrelations across components Global correlations

Considering inter-experiment relationships (Table 2), the most inter-related components are on one hand between BM and IR and the number of chews/bite (r = 0.75) and on the other hand, to a lesser extent, between GT and DMI (r = 0.328). Considering the intra-relationships (Table 2), there are two pairs of variables correlated with each other, independently of BM, on the one hand inter and intra negative relations between BR and chews/bite and on the other hand positive inter and intra relationships between IR and DMI.

**Table 2.** Correlations between ingestive behaviour components implied in the DMI, calculated inter- and intra-experiment (respectively the first and second value per component)

|  | BM | BR | Chews/bite | IR | GT |
| --- | --- | --- | --- | --- | --- |
| **BR** | -0.503***<br>N = 170 |  |  |  |  |
|  | -0.576***<br>N = 386 |  |  |  |  |
| **Chews/bite** | 0.707***<br>N = 43 | -0.597***<br>N = 43 |  |  |  |
|  | 0.760***<br>N = 100 | -0.701***<br>N = 100 |  |  |  |
| **IR**<br>**(mg/min/kg BW)** | 0.703***<br>N = 134 | 0.165**<br>N = 134 | 0.100<sup>ns</sup><br>N = 43 |  |  |
|  | 0.778***<br>N = 313 | -0.244***<br>N = 313 | 0.497***<br>N = 100 |  |  |
| **GT (min/day)** | -0.283**<br>N = 94 | -0.039<sup>ns</sup><br>N = 94 | -0.463*<br>N = 43 | -0.188*<br>N = 94 |  |
|  | -0.117<sup>ns</sup><br>N = 180 | -0.045<sup>ns</sup><br>N = 180 | 0.160<sup>ns</sup><br>N = 100 | -0.239**<br>N = 180 |  |
| **DMI %BW** | 0.243***<br>N = 86 | -0.040<sup>ns</sup><br>N = 86 | 0.645***<br>N = 43 | 0.360***<br>N = 86 | 0.408***<br>N = 86 |
|  | 0.648***<br>N = 153 | -0.254***<br>N = 153 | -0.378<sup>ns</sup><br>N = 100 | 0.793***<br>N = 153 | 0.177*<br>N = 153 |

N = number of data; BM = bite mass; BR = bite rate; IR = intake rate; GT = grazing time.

### Influence of animal species on the relationships

Whatever the type of experience, there is a negative relationship between BR and BM (Figure 5). For cattle, the intra-experiment regression is:

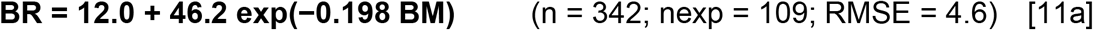

**Figure 5.**
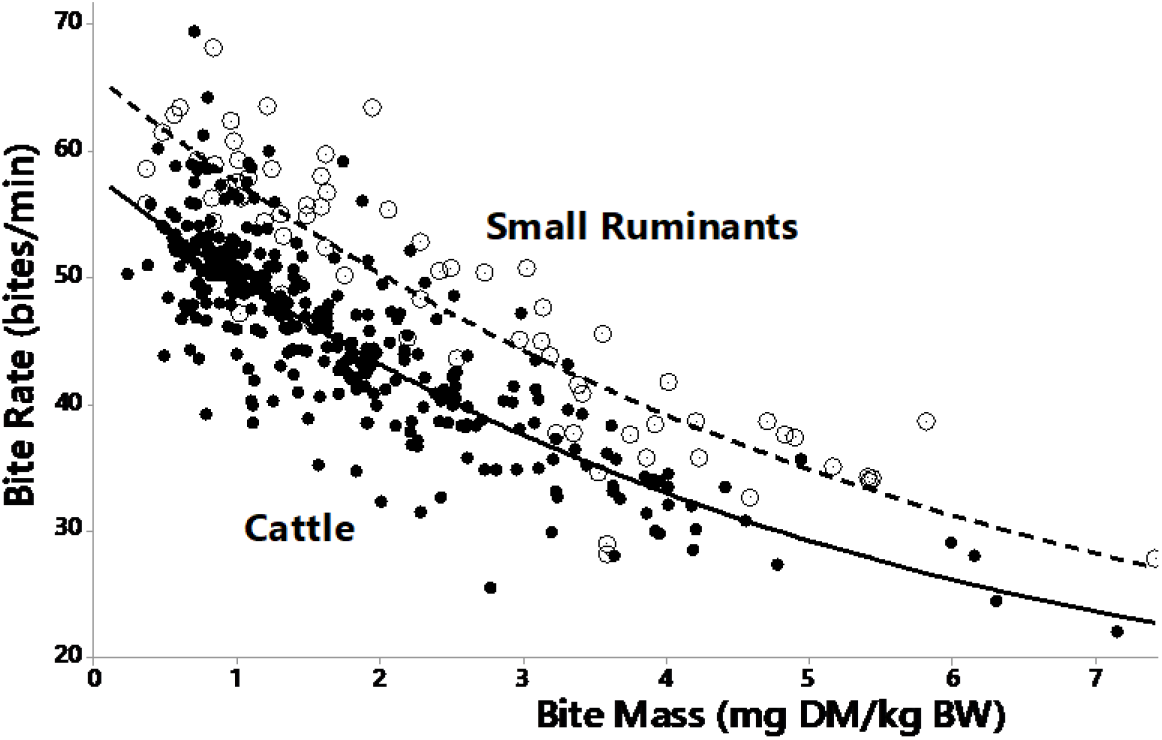
Intra-experiment relationship between bite rate (bites/min) and bite mass (mg/kg BW) for cattle (closed circles) and small ruminants (open circles).

For small ruminants, it is:

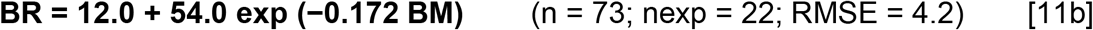

It appears that the asymptote of 12.0 that is never achieved is not different between the two species, while the intercept is significantly higher for small ruminants compared to cattle (66.0 vs 58.2 bites/min).

The JM and chewing associated with the bites are both positively related to BM (Table 2). The relationship between JM/bite and BM is significantly different for small ruminants and cattle (Figure 6a). The intra-species and intra-experiment regression equation for cattle is:

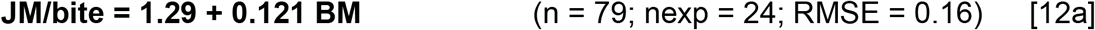

**Figure 6.**
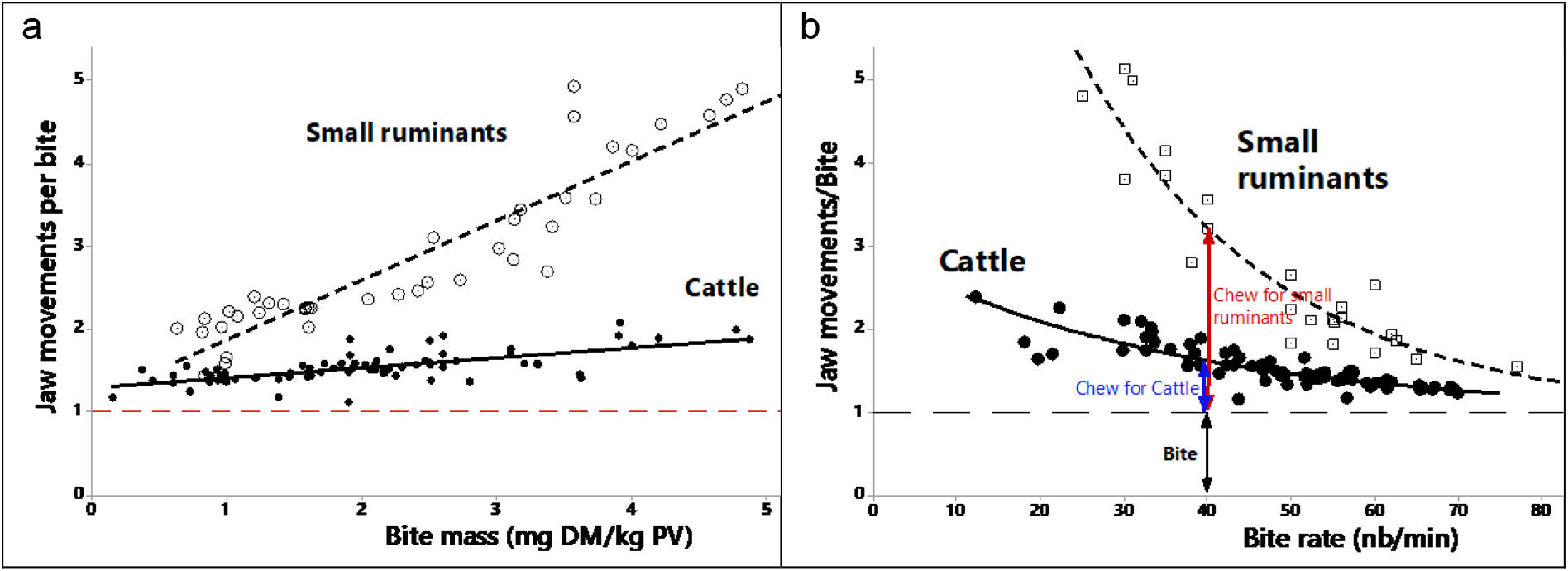
Influences on jaw movements/bite of bite mass (a) and bite rate (b).

For sheep and goats, the corresponding regression is less accurate, and the data number is lower:

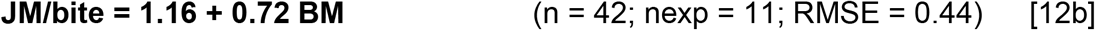

For these two equations, the intercept is not different to 1, illustrating that the number of chews is negligible for very small bites, and in this extreme situation JM are only bites. The data available on jaw and chewing movements also revealed different slopes of the BR-dependent decrease, for cattle and small ruminants, respectively (Figure 6b). For cattle, the intra-experiment regression is:

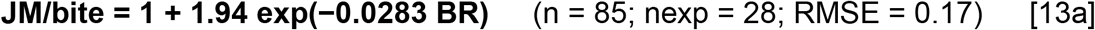

For sheep and goats, the corresponding regression is less accurate, and the data number is lower:

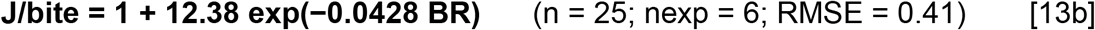

In Figure 6b, the number of JM/bite is the sum of bite + chews per bite. For instance, for a BR of 40/min, the number of JM is about 1.5 JM/bite for cattle, meaning that an animal makes a mean of half a chew/bite. In contrast, for sheep and goats, there are about 2 chews/bite when BR = 40.

It appears clearly that for the same BM, the number of JM is much higher for small ruminants, with an order of magnitude of about 10 (3.1 ± 28.5 vs. 3.3 ± 3.6). The link between JM, expressed per gram of DMI, and BM was also analysed. For both species, the relationship is hyperbolic (Figure 7); the intra-experiment equation for cattle is:

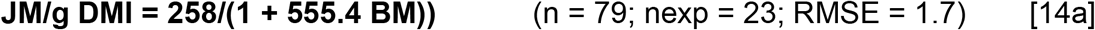

**Figure 7.**
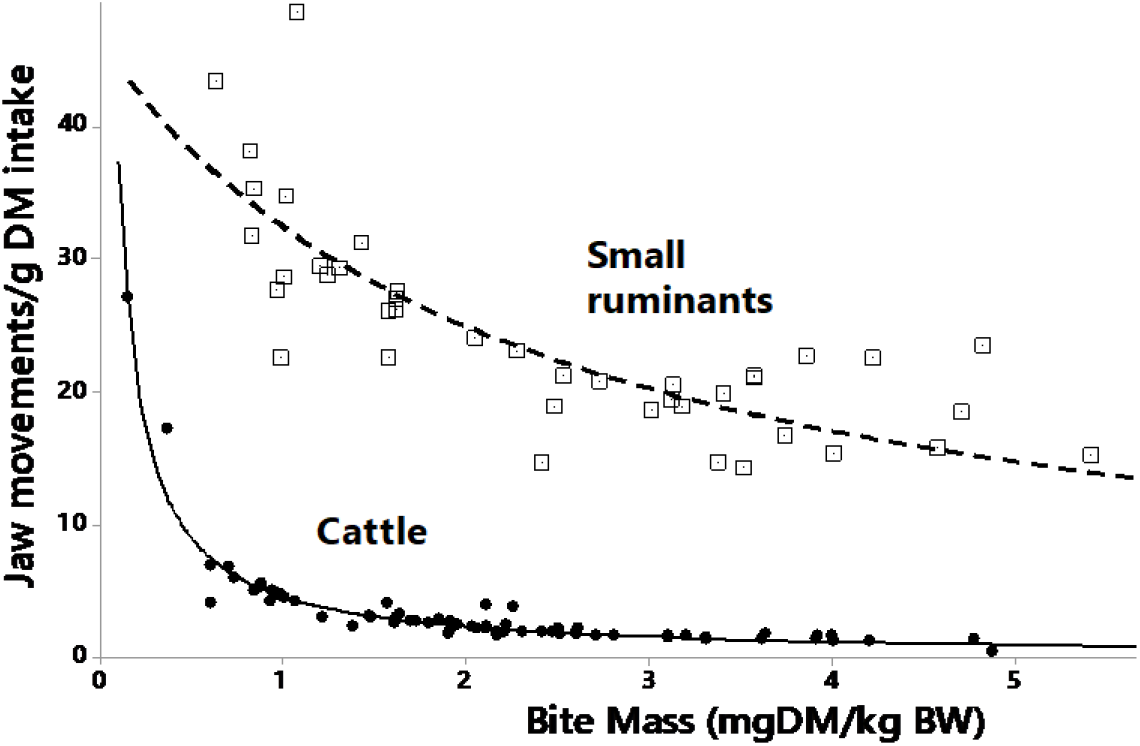
Influence of bite mass (BM, mg DM/kg BW) and of species on the number of jaw movements per gram of DM.

For small ruminants, the regression is less accurate:

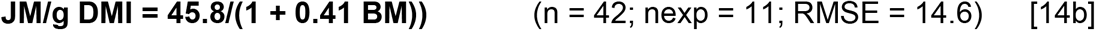

Figure 7 shows these two regressions; it appears clearly that for the same BM, the number of JM is much higher for sheep, with an order of magnitude of about 10 (31.1 ± 28.5 vs 3.3 ± 3.6 JM/g DMI). It must be stressed that for sheep, three high outlier values of 94 to 165 JM/g DMI for a very low BM (BM < 0.4 mg/kg BW) from the same paper (Black and Kenney, 1984) have been removed. Moreover, we were also able to verify the decrease in JM with IR and significant differences that remain between small ruminants and cattle.

As the JM/g DMI are linked to the process of particle comminution, the link between BM and rumination time was investigated from a limited set of data for cattle with 0.5 < BM < 2.5 mg DM/kg BW. It appears that the two components are positively related, according to the following intra-experiment regression:

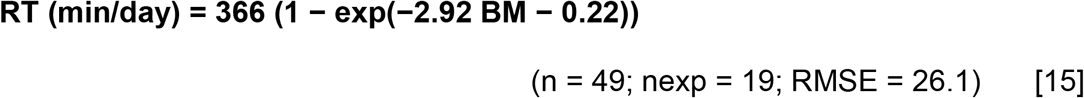

This equation shows an asymptotic value of RT of 366 min/day, and RT drops markedly when BM decreases below a threshold of 1–1.5 mg DM/kg BW. It appears thus that there is a substitution in the comminution activities between intake and rumination. When the fill effect of forage in the mouth increases, the ruminant is less efficient in reducing the particle size so it must ruminate more to compensate.

The relationship between IR (mg DM/min/kg BW) and BM (mg DM/kg BW) is positive and curvilinear and, as BR was different between the two species (Figure 7), two separate fittings were performed. For cattle, the intra-experiment regression is:

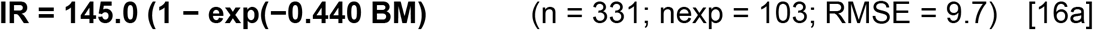

For small ruminants, it is:

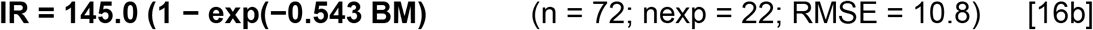

Figure 8 shows the two regressions; it appears that the differences are globally small and are at the advantage of small ruminants for lighter bites, consistent with what was observed for BR (Equations 11a and 11b and Figure 5). The asymptotic value of 145 g DM/kg BW/min is the same between both species. The maximum difference between IR for the two species is observed for BM ~ 2.5 g/kg BW. It must be noted that in order to have a common regression, considering all the data, the power of BW must be 0.85. The curvilinearity of this relationship illustrates that BR, which is the ratio of IR to BM in Figure 5, decreases with the rise of BM as already remarked. Thus, BR is 54 bites/min when BM is close to 0, to approximately 22–23 bites/min when BM is equal to 6 mg/kg BW. This relationship is mainly the outcome of influences of both SH and HBD on BM (Boval and Sauvant, 2019), and IR (Figure 2a and 2b).

**Figure 8.**
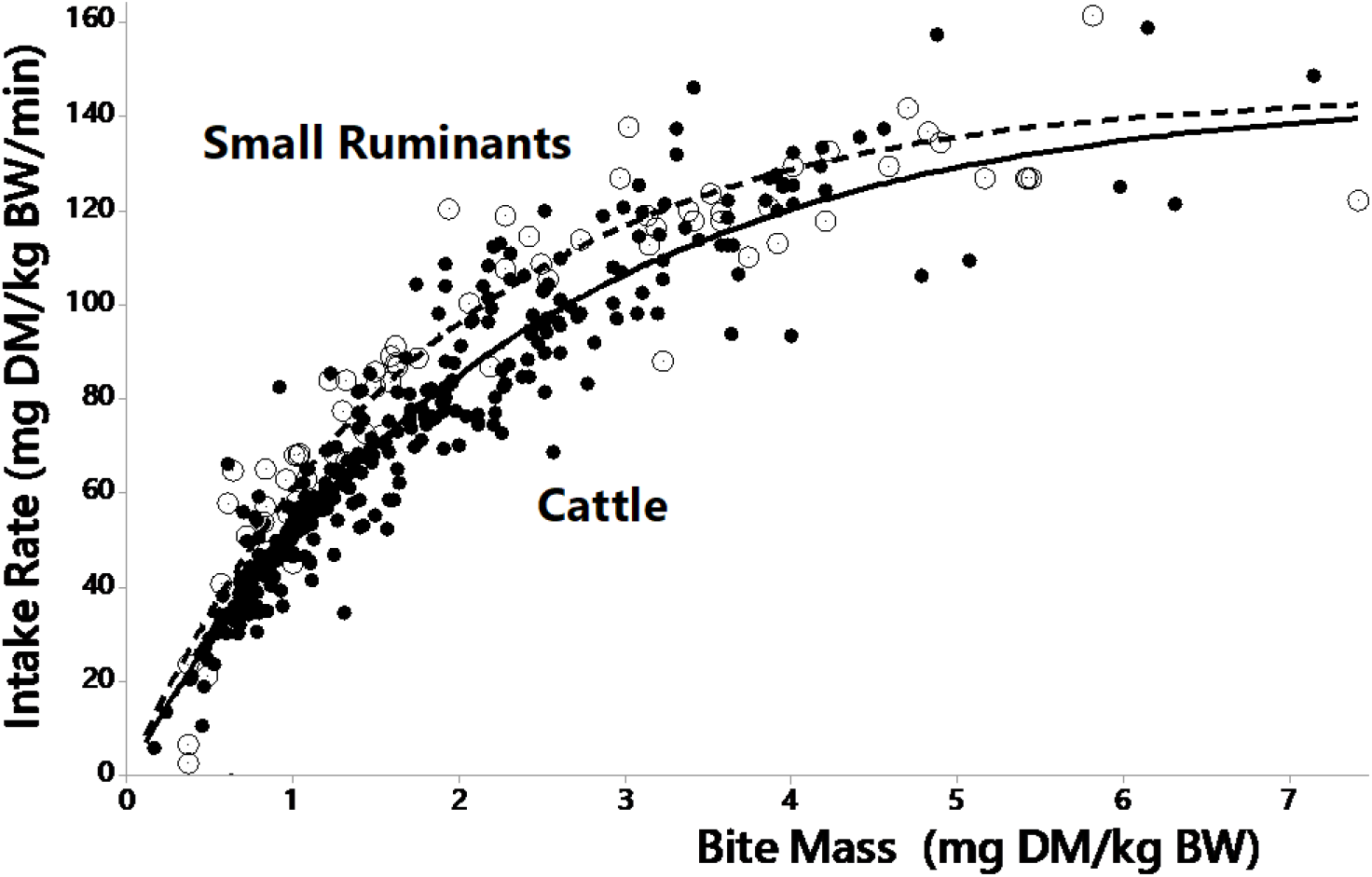
Relationship between intake rate (mg/kg BW/min) and bite mass (mg/kg BW) for cattle (closed circles) and small ruminants (open circles).

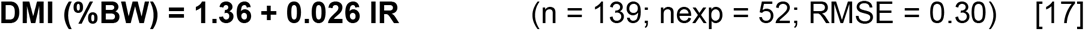

The relationship is still significant when only the 40 experiments focused on the impact of SH are considered. But, in this case the slope is higher i.e. 0.037 (n = 37; nexp = 14; RMSE = 0.13).

There is a positive and curvilinear relationship between BM and daily DMI when treatments with an observation time longer than 1 h are pooled.

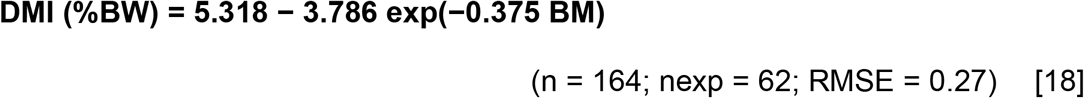

Clearly, a high BM induces a saturated response of both IR (Figure 8) and daily DMI (Figure 9b) in grazing ruminants However, the number of data determining the asymptote is low.

**Figure 9.**
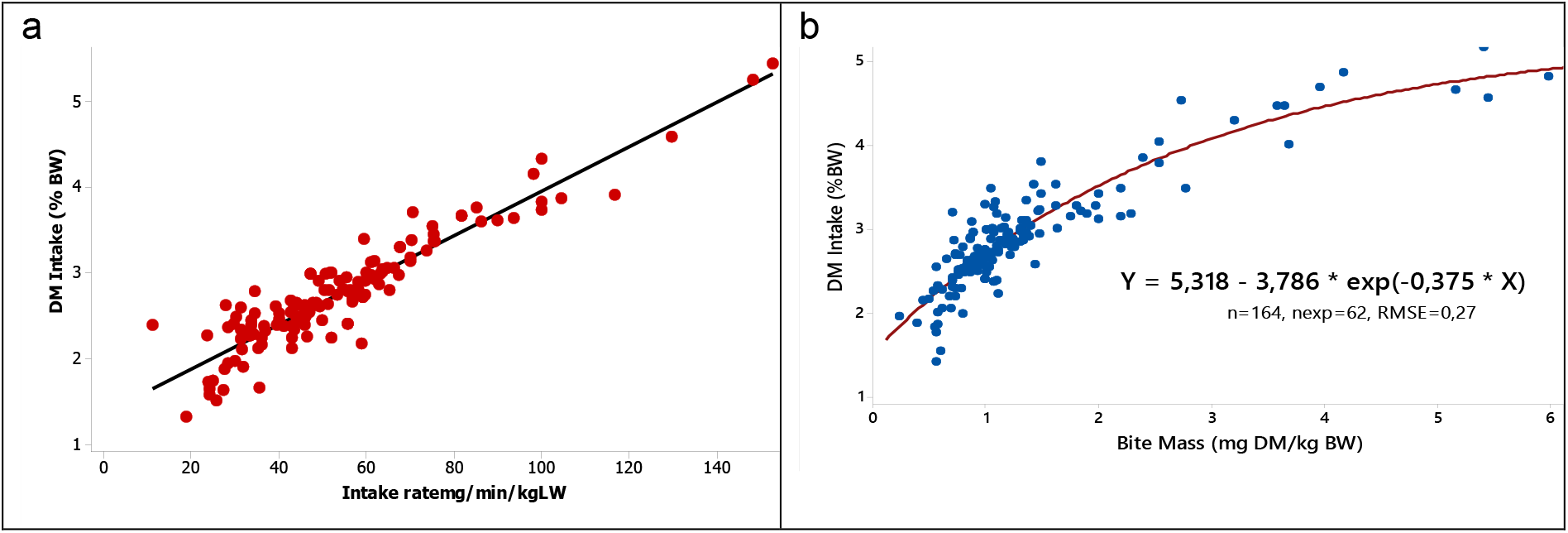
Relationships between daily DM intake (%BW) and intake rate (a) and between bite mass and dry matter intake (b).

## Discussion

### Features of the database

The database made up of 98 publications shows how studies implying cattle predominate, while lines of the database related to small ruminants represent only 1/5 of the total. The publications referenced in this database are spread over the last 40 years, since 1978, with nearly 85% of publications over the last 20 years. The most studied factors of variation in the publications were SH (61 % of the treatments), then bulk density to a much lesser extent (14 % of the treatments). The other factors studied represent less treatments, and the corresponding IB components available were too scattered to allow a valuable interpretation.

Owing to the available data, this meta-analysis presents several limits such as the lack of specific consideration of the impact of some sward characteristics such as the ratio of leaves/stem and their tensile strength or the chemical composition of the sward on the behaviour components. Moreover, we did not consider the spatio-temporal behaviour characteristics of grazing ruminants (feeding stations, patches), nor the kinetics of grazing behaviour during the nycthemeron. Therefore, the considerations done on the time budget are static.

### Impacts of canopy characteristics and of some management strategies

Among the most conventional sward characteristics considered in the literature, SH and HBD mainly have been considered for their impact on BR, IR, GT and DMI. For the other sward characteristics, the IB components available were too scattered to allow a valuable interpretation. Despite the limited data available for other sward characteristics, the effects of herbage mass, LM and SM could have been studied, but only for IR. Moreover, some data were sufficient to be considered under the angle of management strategies, such as HA and access time.

Hence, by increasing SH, BR decreases to a minimum plateau value of about 40 bites/min as soon as the height reaches 20–30 cm (Figure 1a). On the contrary, IR increased with SH, as was previously reported for BM (Boval and Sauvant, 2019), and reached very rapidly a maximum plateau of about 100 mg DM/kg BW/min beyond a height of 20–30 cm (Figure 2a). This maximum plateau results from the combination of the minimum value of BR with the maximum value of BM as proposed by Boval and Sauvant (2019). These trends of response of IR to SH have already been described in the specific contexts of several experiments (Penning, 1986; Ginane and Petit, 2005; Hirata *et al*., 2010). Notably, Delagarde *et al*. (2011) presented a synthetic response of IR with a plateau value close to that of Figure 2a; however, the response was fitted by two linear segments, and presents an elbow that we did not observe at similar values of SH of 22–23 cm. Moreover, for very low values of SH, the decrease of IR was not sufficient compared to their observed data reported in the publication (Delagarde *et al*., 2011), nor to our model. Otherwise, the average plateau calculated in this meta-analysis (Figure 2a) did not include data of three publications (Black and Kenney, 1984; Mezzalira *et al*., 2014 and 2017), where a decrease of IR was observed beyond SH values ranging from 10 to 30 cm (Figure S1). For these same studies, a similar tendency has already been observed for BM (Boval and Sauvant, 2019), suggesting that in certain situations of high SH, it would become more and more difficult to assemble forage into a bite. This consistent decline in IR and BM is thought to be due to the change in the structure of tall species described by some authors (Spallinger and Hobbs, 1992; Mezzalira *et al*., 2014 and 2017).

Regarding the influence of HS on GT, it decreased to a minimum plateau value of about 450 min when SH values exceeded 20–30 cm; while for lower HS values, GT becomes higher and can exceed 650 min/day (Figure S3). This trend is consistent with previous reports by Alvarez *et al*. (2007) and Perez-Prieto *et al*. (2011).

By increasing the HBD, BR decreased until a plateau of around 40 bites/min beyond 2–3 kg DM/m^3^ (Figure 1b), similar to the minimum plateau observed with high SH (Figure 1a) in another set of experiments. We noted a similar trend with IR, which also decreased globally with HBD (Figure 2b) and plateaued at about 60 mg DM/kg BW/min, at the same HBD threshold of 2–3 kg DM/m^3^. These trends are globally opposite to what were previously observed for BM, which increases with both SH and HBD (Boval and Sauvant, 2019). Unfortunately, concerning GT, the effect of HBD could not be analysed as for SH, as most of the experiments that tested HBD variations were carried out with micro-swards, for which the duration of grazing could not be measured.

In fact, the effects of SH and HBD cannot be disconnected from one another in natural grazing conditions, and we were opportunely able to study in our database the interaction between these two major factors, on IR (Figure 3). Globally the effect of SH is more marked than that of HBD. At low SH, HBD positively influences IR while for high SH, HBD presents a limited negative influence on IR (Figure 3). Only a few individual studies have measured this interaction between SH and HBD consistently to our results. It has been studied actually only for short grasses and with micro-swards, as by Laca *et al*. (1992) and Benvenutti *et al*. (2006).

Aside from SH and bulk density, other characteristics are also likely to influence IR, such as LM and SM. They both have a positive effect on IR; in particular, LM explains a larger range of IR, surely linked with leaf growth, without any threshold being observed. In contrast, when the SM increases beyond 1 t of DM/ha, the IR reaches a plateau of 100 mg MS/kg BW/min.

While our database was not mainly focused on the influence of management strategies on global responses such as DMI, some publications allowed highlighting of some IB responses. Thus, ruminants are able to increase their IR until values near to 100 mg DMI/kg BW/min when facing an important decrease of forage allowance (Figure 4a) or of grazing access time (Figure 4b). This adaptive behaviour mainly results from differences in BM (Boval and Sauvant, 2019) which appears as a key factor of animal robustness as it allows ruminants to maintain, or only slightly decrease, their level of DMI despite a decrease of available resource and access time. As BM is at least partly explained by individual factors (Sollenberger and Vanzant, 2011; Boval and Sauvant, 2019), it appears useful to investigate further animals’ ability to adapt to restricted resources and GT.

### From BM to intake rate and daily intake

Analysis of the impact of canopy characteristics on the various IB components highlights the behavioural adaptation by ruminants to achieve satisfying BM and IR. That appears for the low values of SH (< 20–30 cm) and of HBD (< 2–3 kg DM/m^3^), resulting in an acceleration of BR (Figure 1a and 1b) to compensate for the smaller bites. Consequently, the resulting IR is actually increased at low HBD (Figure 2b), while that is not the case at low SH (Figure 2a) due to the first limiting effect of SH on BM (Boval and Sauvant, 2019). In addition, for low SH, the GT is longer (Figure S3), as another way to compensate for low values of BM and IR. However, we did not have enough data to show this lengthening of GT also with low values of HBD.

Beyond analysis of the effect of sward characteristics on IB components, the major relationships between these components provided further understanding. Thus, there is a strong negative correlation between BM and BR (Table 2); correlations between BM and chews/bite are also positive, while based on much less data (Table 2), and the correlation between BM and DMI is less marked, especially inter-experiment correlation. And most structuring regressions concern the link between BM and BR and the influence of BM on IR (Figure 8) which are useful for modelling purposes. All these correlations are consistent with previous reports (Poppi, 2011; Chilibroste *et al*., 2007 and 2015) and this meta-analysis, resulting from numerous data, provides robust average values of the main correlations.

The close negative relationship between BM and BR may be better understood by analysing JM/bite (Figure 6a) and how they increase with BM. Grazing animals perform JM, which contribute both to assembling the forage before harvesting and to chewing it in the mouth, before swallowing. Hence, larger bites require logically more processing before the next bite can be taken (Mulvenna *et al*., 2018). Therefore, the time needed between two bites increases, representing the sum of the time devoted in the JM to biting and chewing. These additional activities mechanically slow down the frequency of bites (Figure 6b). According to Spallinger and Hobbs (1992), BR is indeed the inverse of handling time (i.e. the time invested to bite and chew) and this occurs mainly in pastures, where potential bites are concentrated, corresponding mostly to a functional response of type 3, according to Mezzalira *et al*. (2017).

The analysis of JM expressed per gram of DM consumed (Figure 7) shows that the number of JM decreases when the bites become larger. This suggests that with larger bites of more than 1 mg/kg BW, particle fragmentation efficiency decreases (Sauvant *et al*., 1996; Baumont et al, 2000). This is consistent with our results showing how larger bites are positively correlated with longer rumination times (Figure S4, Equation 15). With larger bites, the fill effect of the forage in the mouth increases and the grazer would be less efficient in reducing the particle size and so it must ruminate more to compensate. It appears then as a substitution in the comminution activities between intake and rumination. This could be due also partly to the fact that small BM is more often composed of more fibrous removed parts. Indeed, larger bites are more often associated with the presence of leaves in the sward canopy (Drescher *et al*., 2006; Geremia *et al*., 2018), whereas the more fibrous stems represent a physical resistance inducing a limit to biting. Geremia *et al*. (2018) reported how BM is small at the end of the grazing period, as animals are forced to harvest grass with a higher percentage of stems and dead material.

Clearly, BM is the major determinant of IR and contributes consequently to differences of DMI. Our dataset contains some large BM, more than 4 mg/kg BW, which allows very high values of IR to be achieved, up to 140 mg/kg BW (Figure 8), while the mean asymptotic values observed for factors were around 100 mg/kg BW (Figures 2–4). Several authors had already reported these positive relationships (ref), and our results provide few data for extreme situations (low SH and HBD), as 140 mg appears as a maximum rate achievable whatever the animal species. For these few high BM and IR values, the corresponding DM intake may exceed 5% of BW, being also influenced by the total daily duration of grazing. However, contrary to the curvilinear relationship between BM and DMI (Figure 9b), the relationship between IR and DMI (Figure 9a) appears linear, and we could not highlight any threshold of DMI. These results could be due to the low number of DMI values corresponding to high values of IR that we collected.

### Animal species specificity

From our database, we were able to calculate some specific relationships for cattle or small ruminants, being unable to distinguish between sheep and goats. However, according to Mulvena *et al*. (2018) and Laca (2010), there is no marked difference between these two species of small ruminants.

It appears that for the same level of BM, small ruminants graze with a faster BR compared to cattle, by about 10 bites/min (Figure 5). Besides that, small ruminants make more JM accompanying each bite compared to cattle (Figure 6b), and the difference increases clearly when BR decreases. For very small bites less than 1 mg/kg BW, there are almost no JM/bite (Figure 6a). For small ruminants, the number of chews increases very quickly, with BM of 5 mg/kg BW, requiring about 5 JM/bite (Figure 6a). Thus when JM are expressed per gram of DMI, the number of JM is approximately 10 times higher for small ruminants compared with cattle (Figure 7). It is approximately the same scaling value when both species are compared in terms of DMI. In any case, small ruminants make many more JM to crop 1 g of DMI, likely investing more energy per gram of DMI than cattle, as already reported (Galli *et al*., 2018). Aside from chews, another type of JM may explain the difference between species, i.e. the chew-bite that can be measured with some acoustic monitoring methods (Galli *et al*., 2018). Indeed, cattle (Ungar *et al*., 2006) and sheep (Galli *et al*., 2011) may use discrete JM to chew and bite, but also simultaneously chew and bite on the same jaw opening-closing cycle. However, in our database, we had no such values of chew-bites.

Consequently, all of these differences imply that small ruminants have a faster IR, with a maximum equal to 1.2 times higher IR expressed per kg of BW, compared to cattle (Figure 8). Otherwise, Boval and Sauvant (2019) have also pointed out that, for the same SH, the bite depth/kg BW is higher for small ruminants, so it was necessary to use BW at power 0.20 to match the data of BD for both species. The difference of bite depth between the two species is extremely low compared to their respective BW, revealing that sheep chewing and biting modalities would be more effective for going deeper into the sward compared to cattle, as already suggested (Gordon *et al*., 1996; Woodward, 1998; Baumont *et al*., 2006). Indeed, it may be observed how sheep perform successive chews to go deeper into the canopy sward, by mobilizing their lips in quick movements.

All these results are well consistent with the idea that bite and chew rates decrease commonly with ruminant species having greater BW and BM (Wilson and Kerley, 2003; Mulvenna *et al*., 2018). Moreover, these results emphasize the different mechanisms implemented by small ruminants or cattle to adapt to the characteristics of the resource, with their anatomical specificities (Baumont *et al*., 2006; Meier *et al*., 2016). In the case of small ruminants, the most mobile lips participate in forage prehension and therefore consumption, with associated movements of the jaw that can be recorded. Cattle have a particularly long freely mobile tip (Meier *et al*., 2016) that they use to greatly increase the diameter and surface of each bite to compensate for limited resources as with short SH (Boval and Sauvant, 2019). Although the relationships between the different behavioural variables are different for these two species, the fact remains that the make-up of the bite for both species determines the rate of intake with very similar trends (Figure 9) and with a comparable maximum threshold around 140 mg/kg BW).

## Conclusions

This meta-analysis provided a set of empirical models that can serve (i) as benchmarks for future studies and models of ruminant feeding behavior and well-being (ii) to identify parameters of interest for animal management at pasture (iii) to reference values for automatic measurement devices.

Approximately 20 quantitative relationships were established within this meta-analysis, confirming that bite size is a pivotal part of the ingestive behavior of ruminants in pasture, as it is both sensitive to major sward characteristics and determining for intake rate, and daily intake.

The main response laws highlighted are valid for different domestic ruminants when expressed in relation to body weight. Nevertheless, important differences appeared between cattle and small ruminants, the latter having to chew more for the same bite mass. The literature review emphasizes the great variability of methods carried out to measure ingestive behavior components. Our database should be supplemented by data collected with animals in stalls to assess the generic relationships applying whatever the feeding context.

## Supplementary material

**Figure S1:**
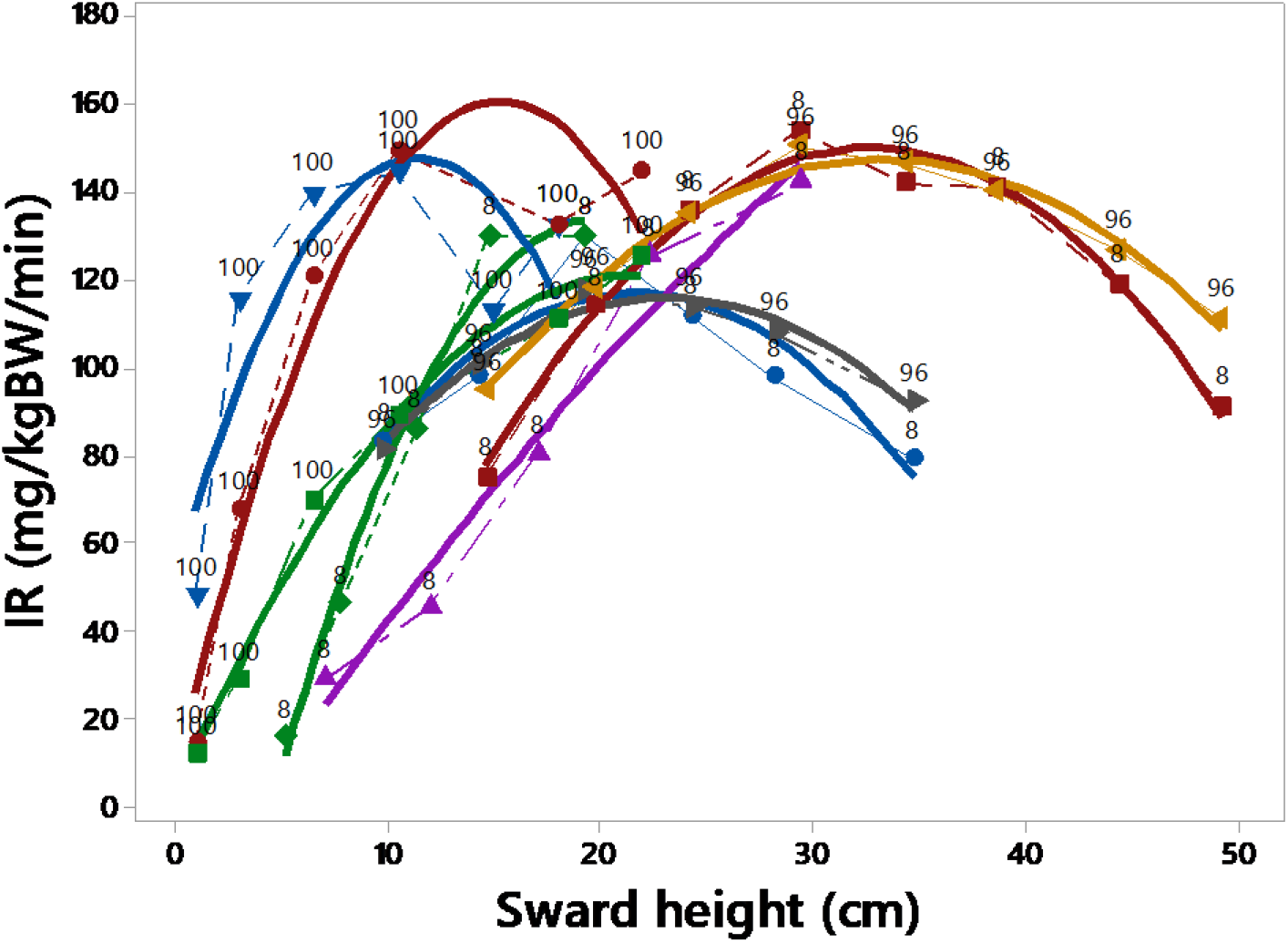
Impact of sward height on Intake rate for some studies

**Figure S2:**
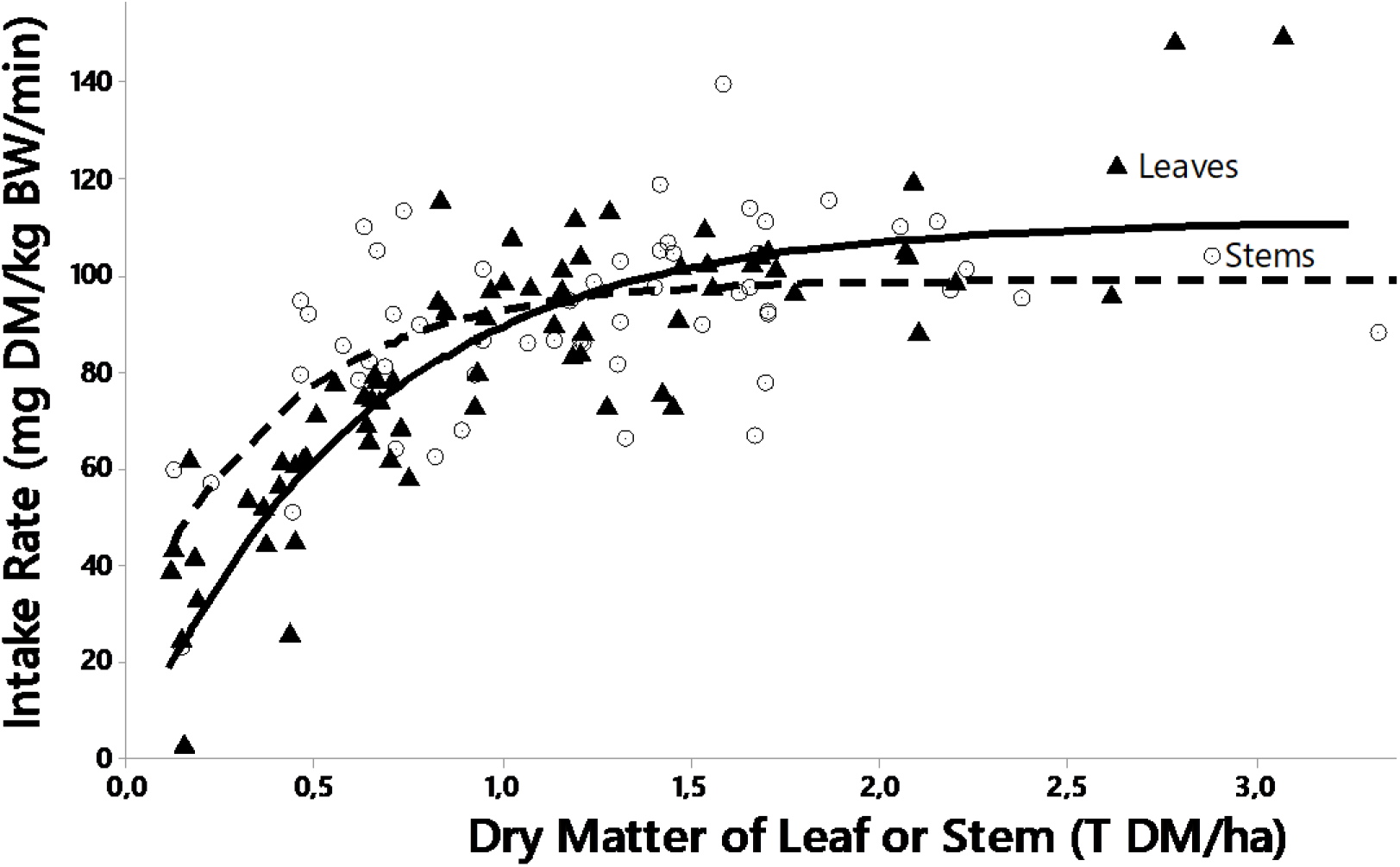
Respective influences of DM of leaf and stem, on intake rate.

**Figure S3:**
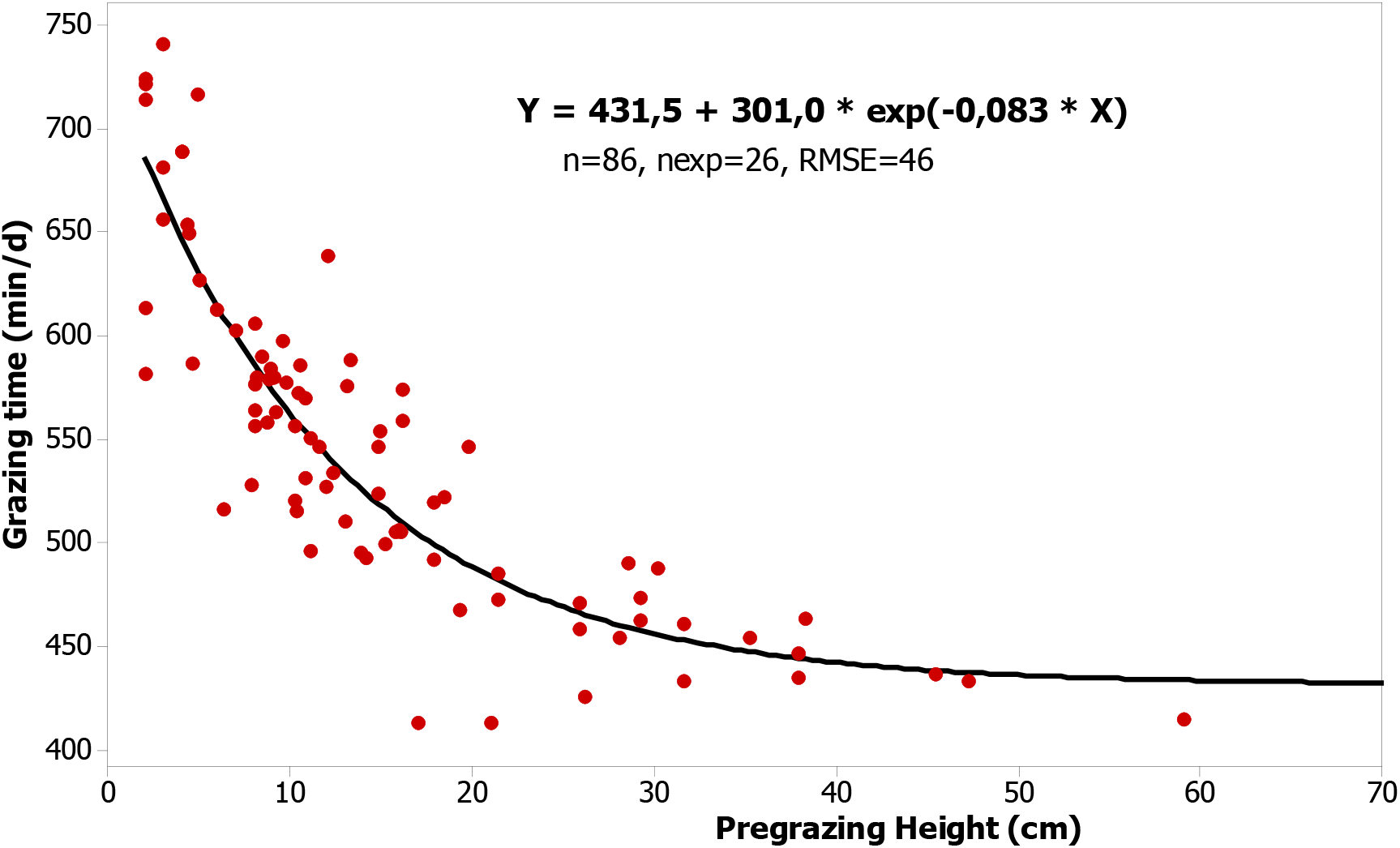
Effect of sward height (cm) on grazing time (min/day).

**Figure S4:**
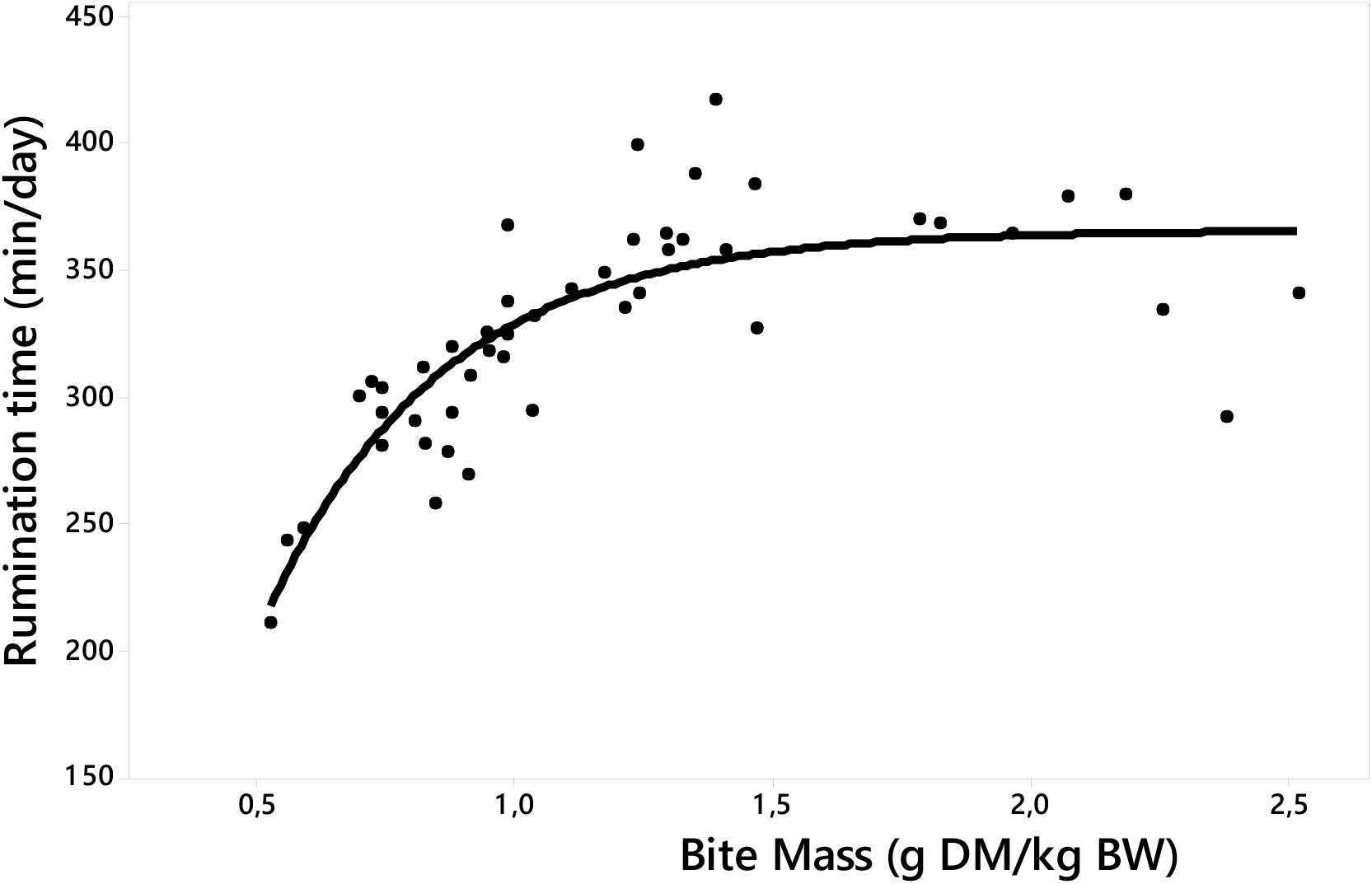
Relationship between rumination time (min/day) and Bite mass (mg/kg BW)

## References used for the meta-analysis

Abrahamse PA, Tamminga S and Dijkstra J 2009. Effect of daily movement of dairy cattle to fresh grass in morning or afternoon on intake, grazing behaviour, rumen fermentation and milk production. Journal of Agricultural Science 147, 721–730.

Ackerman CJ, Purvis HT, Horn GW, Paisley SI, Reuter RR and Bodine TN 2001. Performance of light vs heavy steers grazing Plains Old World bluestem at three stocking rates. Journal of Animal Science 79, 493–499.

Amaral MF, Mezzalira JC, Bremm C, Da Trindade JK, Gibb MJ, Sune RWM and Carvalho PCD 2013. Sward structure management for a maximum short-term intake rate in annual ryegrass. Grass and Forage Science 68, 271–277.

Ayantunde AA, Fernandez-Rivera S, Hiernaux PH and Tabo R 2008. Implications of restricted access to grazing by cattle in wet season in the Sahel. Journal of Arid Environments 72, 523–533.

Ayantunde AA, Fernandez-Rivera S, Hiernaux PHY, Keulen Hv, Udo HMJ and Chanono M 2001. Effect of timing and duration of grazing of growing cattle in the West African Sahel on diet selection, faecal output, eating time, forage intake and live-weight changes. Animal Science 72, 117–128.

Barbieri CW, de Quadros FLF, Jochims F, Kuinchtner BC, de Carvalho THN, Casanova PT, Fernandes AM and Pereira JB 2015. Beef heifers grazing behavior and herbage intake in natural grassland under rotational grazing. Ciencia Rural 45, 2056–2062.

Barrett PD, McGilloway DA, Laidlaw AS and Mayne CS 2003. The effect of sward structure as influenced by ryegrass genotype on bite dimensions and short-term intake rate by dairy cows. Grass and Forage Science 58, 2–11.

Benvenutti MA, Gordon IJ and Poppi DP 2006. The effect of the density and physical properties of grass stems on the foraging behaviour and instantaneous intake rate by cattle grazing an artificial reproductive tropical sward. Grass and Forage Science 61, 272–281.

Benvenutti MA, Gordon IJ and Poppi DP 2008. The effects of stem density of tropical swards and age of grazing cattle on their foraging behaviour. Grass and Forage Science 63, 1–8.

Benvenutti MA, Gordon IJ, Poppi DP, Crowther R, Spinks W and Moreno FC 2009. The horizontal barrier effect of stems on the foraging behaviour of cattle grazing five tropical grasses. Livestock Science 126, 229–238.

Black JL and Kenney PA 1984. Factors affecting diet selection by sheep .2. Height and density of pasture. Australian Journal of Agricultural Research 35, 565–578.

Bodine TN and Purvis HT, II 2003. Effects of supplemental energy and/or degradable intake protein on performance, grazing behavior, intake, digestibility, and fecal and blood indices by beef steers grazed on dormant native tallgrass prairie. In, pp. 304–317.

Boval M, Archimède H, Cruz P and Duru M 2007a. Intake and digestibility in a Dichanthium spp. dominated pasture, at 14 an 28 days of regrowth. Animal Feed Science and Technology 134, 18–31.

Boval M, Archimede H, Tournebize R and Coppry O 2002a. Stage of regrowth of tropical forage have various effect on diet quality of grazing heifers. In 19th General Meeting of European Grassland Federation, pp. 108–109. La Rochelle, France.

Boval M, Cruz P, Ledet JE, Coppry O and Archimede H 2002b. Effect of nitrogen on intake and digestibility of a tropical grass grazed by Creole heifers. Journal of Agricultural Science 138, 73–84.

Boval M, Cruz P, Peyraud JL and Penning P 2000. The effect of herbage allowance on daily intake by Creole heifers tethered on natural Dichanthium spp. pasture. Grass and Forage Science 55, 201–208.

Boval M, Fanchone A, Archimede H and Gibb MJ 2007b. Effect of structure of a tropical pasture on ingestive behaviour, digestibility of diet and daily intake by grazing cattle. Grass and Forage Science 62, 44–54.

Braghieri A, Pacelli C, Girolami A and Napolitano F 2011. Time budget, social and ingestive behaviours expressed by native beef cows in Mediterranean conditions. Livestock Science 141, 47–52.

Bratti LFS, Dittrich JR, Barros CS, Silva CJA, Monteiro ALG, Rocha CDR, Rocha FMPD 2009. Ingestive behavior of goats in ryegrass and black oat pastures in pure or mixture culture. Ciência Animal Brasileira, 10(2), 397–405.

Bremm C, Laca EA, Fonseca L, Mezzalira JC, Elejalde DAG, Gonda HL and Carvalho PCD 2012. Foraging behaviour of beef heifers and ewes in natural grasslands with distinct proportions of tussocks. Applied Animal Behaviour Science 141, 108–116.

Brink GE and Soder KJ 2011. Relationship between Herbage Intake and Sward Structure of Grazed Temperate Grasses. Crop Science 51, 2289–2298.

Burlison AJ, Hodgson J and Illius AW 1991. Sward canopy structure and the bite dimensions and bite weight of grazing sheep. Grass and Forage Science 46, 29–38.

Cangiano CA, Galli JR, Pece MA, Dichio L and Rozsypalek SH 2002. Effect of liveweight and pasture height on cattle bite dimensions during progressive defoliation. Australian Journal of Agricultural Research 53, 541–549.

Casey IA, Laidlaw AS, Brereton AJ, McGilloway DA and Watson S 2004. The effect of bulk density on bite dimensions of cattle grazing microswards in the field. Journal of Agricultural Science 142, 109–121.

Chacon EA, Stobbs TH and Dale MB 1978. Influence of Sward Characteristics on Grazing Behavior and Growth of Hereford Steers Grazing Tropical Grass Pastures. Australian Journal of Agricultural Research 29, 89–102.

Charpentier A and Delagarde R 2018. Milk production and grazing behaviour responses of Alpine dairy goats to daily access time to pasture or to daily pasture allowance on temperate pastures in spring. Small Ruminant Research 162, 48–56.

Costa JV, Oliveira ME, Moura R, da Costa MDN and Rodrigues MM 2015. Grazing behavior and ingestive goats in silvopastoral system. Revista Ciencia Agronomica 46, 865–872.

Da Trindade JK, Neves FP, Pinto CE, Bremm C, Mezzalira JC, Nadin LB, Genro TCM, Gonda HL and Carvalho PCF 2016. Daily Forage Intake by Cattle on Natural Grassland: Response to Forage Allowance and Sward Structure. Rangeland Ecology & Management 69, 59–67.

Da Trindade JK, Pinto CE, Neves FP, Mezzalira JC, Bremm C, Genro TCM, Tischler MR, Nabinger C, Gonda HL and Carvalho PCF 2012. Forage Allowance as a Target of Grazing Management: Implications on Grazing Time and Forage Searching. Rangeland Ecology & Management 65, 382–393.

de Macedo JDB, Teixeira FA, da Silva FF, Pires AJV, de Aguilar PB and Nascimento PVN 2016. Ingestive behavior of heifers on Brachiaria pasture deferred with different periods of sealing. Semina-Ciencias Agrarias 37, 4275–4284.

de Souza ANM, da Rocha MG, Potter L, Roso D, Glienke CL and Neto RAD 2011. Ingestive behavior of beef heifers in warm season annual grass pastures. Revista Brasileira De Zootecnia-Brazilian Journal of Animal Science 40, 1662–1670.

Dias DLS, Silva RR, Silva FFd, Carvalho GGPd, Barroso DS and Carvalho VM 2014. Correlation between performance and ingestive behavior of steers post-weaned on pastures. Acta Scientiarum. Animal Sciences 36, 93–99.

Dohme-Meier F, Kaufmann LD, Gors S, Junghans P, Metges CC, van Dorland HA, Bruckmaier RM and Munger A 2014. Comparison of energy expenditure, eating pattern and physical activity of grazing and zero-grazing dairy cows at different time points during lactation. Livestock Science 162, 86–96.

Drescher M, Heitkonig IMA, Van den Brink PJ and Prins HHT 2006. Effects of sward structure on herbivore foraging behaviour in a South African savanna: An investigation of the forage maturation hypothesis. Austral Ecology 31, 76–87.

Dumont B, Garel JP, Ginane C, Decuq F, Farruggia A, Pradel P, Rigolot C and Petit M 2007. Effect of cattle grazing a species-rich mountain pasture under different stocking rates on the dynamics of diet selection and sward structure. Animal 1, 1042–1052.

Fanchone A, Archimede H, Baumont R and Boval M 2010. Intake and digestibility of fresh grass fed to sheep indoors or at pasture, at two herbage allowances. Animal Feed Science and Technology 157, 151–158.

Fanchone A, Archimede H, Delagarde R and Boval M 2012. Comparison of intake and digestibility of fresh Digitaria decumbens grass fed to sheep, indoors or at pasture, at two different stages of regrowth. Animal 6, 1108–1114.

Fernandes M, Fernandes JS, De Resende KT, Bonfa HC, Reis RA, Ruggieri AC, Fernandes JJR and Santos PM 2016. Grazing behavior and intake of goats rotationally grazing Tanzania-grass pasture with different post-grazing residues. Tropical Grasslands-Forrajes Tropicales 4, 91–100.

Flores ER, Laca EA, Griggs TC and Demment MW 1993. Sward height and vertical morphological-differentiation determine cattle bite dimensions. Agronomy Journal 85, 527–532.

Fonseca L, Carvalho PCF, Mezzalira JC, Bremm C, Galli JR and Gregorini P 2013. Effect of sward surface height and level of herbage depletion on bite features of cattle grazing Sorghum bicolor swards. Journal of Animal Science 91, 4357–4365.

Galli JR, Cangiano CA, Milone DH and Laca EA 2011. Acoustic monitoring of short-term ingestive behavior and intake in grazing sheep. Livestock Science 140, 32–41.

Galli JR, Cangiano CA, Pece MA, Larripa MJ, Milone DH, Utsumi SA and Laca EA 2018. Monitoring and assessment of ingestive chewing sounds for prediction of herbage intake rate in grazing cattle. Animal 12, 973–982.

Gibb MJ, Huckle CA and Nuthall R 1998. Effect of time of day on grazing behaviour by lactating dairy cows. Grass and Forage Science 53, 41–46.

Gibb MJ, Huckle CA, Nuthall R and Rook AJ 1997. Effect of sward surface height on intake and grazing behaviour by lactating Holstein Friesian cows. Grass and Forage Science 52, 309–321.

Ginane C, Petit M and D’Hour P 2003. How do grazing heifers choose between maturing reproductive and tall or short vegetative swards? Applied Animal Behaviour Science 83, 15–27.

Glienke CL, Rocha MG, Potter L, Roso D, Montagner DB and Neto RAO 2016. Canopy structure, ingestive behavior and displacement patterns of beef heifers grazing warm-season pastures. Arquivo Brasileiro De Medicina Veterinaria E Zootecnia 68, 457–465.

Goncalves EN, Carvalho PCD, Kunrath TR, Carassai IJ, Bremm C and Fischer V 2009. Plant-animal relationships in pastoral heterogeneous environment: process of herbage intake. Revista Brasileira De Zootecnia-Brazilian Journal of Animal Science 38, 1655–1662.

Goncalves RP, Bremm C, Moojen FG, Marchi D, Zubricki G, Caetano LAM, Neto AB and Carvalho PCD 2018. Grazing down process: The implications of sheep’s ingestive behavior for sward management. Livestock Science 214, 202–208.

Gordon IJ, Illius AW and Milne JD 1996. Sources of variation in the foraging efficiency of grazing ruminants. Functional Ecology 10, 219–226.

Gregorini P, Clark C, McLeod K, Glassey C, Romera A and Jago J 2011. Feeding station behavior of grazing dairy cows in response to restriction of time at pasture. Livestock Science 137, 287–291.

Gregorini P, Gunter SA, Beck PA, Caldwell J, Bowman MT and Coblentz WK 2009. Short-term foraging dynamics of cattle grazing swards with different canopy structures. Journal of Animal Science 87, 3817–3824.

Gregorini P, Gunter SA, Bowman MT, Caldwell JD, Masino CA, Coblentz WK and Beck PA 2011. Effect of herbage depletion on short-term foraging dynamics and diet quality of steers grazing wheat pastures. Journal of Animal Science 89, 3824–3830.

Griffiths WM, Hodgson J and Arnold GC 2003. The influence of sward canopy structure on foraging decisions by grazing cattle. I. Patch selection. Grass and Forage Science 58, 112–124.

Hampel VDS, da Rocha MG, Poetter L, Stivanin SCB, Alves MB, Cado LM, Neto LGD and Vicente JM 2016. Grazing behavior of non-supplemented and supplemented heifers on Italian ryegrass pasture. Semina-Ciencias Agrarias 37, 2053–2065.

Hirata M, Matsumoto Y, Izumi S, Soga Y, Hirota F and Tobisa M 2015. Seasonal and interannual variations in feeding station behavior of cattle: effects of sward and meteorological conditions. Animal 9, 682–690.

Hirata M, Sakou A, Terayama Y, Furuya M and Nanba T 2008. Selection of feeding areas by cattle in a spatially heterogeneous environment: selection between two tropical grasses. Journal of Ethology 26, 327–338.

Hirata M, Yamamoto K and Tobisa M 2010. Selection of feeding areas by cattle in a spatially heterogeneous environment: selection between two tropical grasses differing in accessibility and abiotic environment. Journal of Ethology 28, 95–103.

Idibu J, Kabi F and Mpairwe D 2016. Behavioural response of pure Ankole and crossbred (Ankole x Holstein) cows to seasonal pasture variations in south-western Uganda. Applied Animal Behaviour Science 174, 11–18.

Illius AW, Gordon IJ, Milne JD and Wright W 1995. Costs and benefits of foraging on grasses varying in canopy structure and resistance to defoliation. Functional Ecology 9, 894–903.

Laca EA, Ungar ED and Demment MW 1994. Mechanisms of Handling Time and Intake Rate of a Large Mammalian Grazer. Applied Animal Behaviour Science 39, 3–19.

Laca EA, Ungar ED, Seligman N and Demment MW 1992. Effects of sward height and bulk-density on bite dimensions of cattle grazing homogeneous swards. Grass and Forage Science 47, 91–102.

Lee C, Fisher AD, Colditz IG, Lea JM and Ferguson DM 2013. Preference of beef cattle for feedlot or pasture environments. Applied Animal Behaviour Science 145, 53–59.

Mattiauda DA, Tamminga S, Gibb MJ, Soca P, Bentancur O and Chilibroste P 2013. Restricting access time at pasture and time of grazing allocation for Holstein dairy cows: Ingestive behaviour, dry matter intake and milk production. Livestock Science 152, 53–62.

McCarthy S, Horan B, Rath M, Linnane M, O’Connor P and Dillon P 2007. The influence of strain of Holstein-Friesian dairy cow and pasture-based feeding system on grazing behaviour, intake and milk production. Grass and Forage Science 62, 13–26.

McGilloway DA, Cushnahan A, Laidlaw AS, Mayne CS and Kilpatrick DJ 1999. The relationship between level of sward height reduction in a rotationally grazed sward and short-term intake rates of dairy cows. Grass and Forage Science 54, 116–126.

Mendes FBL, Silva RR, de Carvalho GGP, da Silva FF, Lins T, da Silva ALN, Macedo V, Abreu G, de Souza SO and Guimaraes JO 2015. Ingestive behavior of grazing steers fed increasing levels of concentrate supplementation with different crude protein contents. Tropical Animal Health and Production 47, 423–428.

Mezzalira JC, Bonnet OJF, Carvalho PCD, Fonseca L, Bremm C, Mezzalira CC and Laca EA 2017. Mechanisms and implications of a type IV functional response for short-term intake rate of dry matter in large mammalian herbivores. Journal of Animal Ecology 86, 1159–1168.

Mezzalira JC, Carvalho PCD, Fonseca L, Bremm C, Cangiano C, Gonda HL and Laca EA 2014. Behavioural mechanisms of intake rate by heifers grazing swards of contrasting structures. Applied Animal Behaviour Science 153, 1–9.

Mitchell RJ, Hodgson J and Clark DA 1991. The effect of varying leafy sward height and bulk-density on the ingestive behavior of young deer and sheep. In Proceedings of the New Zealand Society of Animal Production, Vol 51 1991 (ed. A Parry), pp. 159–165.

Moterle PH, Rocha MG, Potter L, Sichonany MJO, Neto L, Silva MF, Salvador PR and Vicente JM 2017. Displacement patterns of beef heifers receiving supplement in Italian ryegrass pasture. Arquivo Brasileiro De Medicina Veterinaria E Zootecnia 69, 1021–1029.

Neto RAD, da Silva JHS, da Rocha MG, Potter L, Sichonany MJD, Biscaino LL, dos Santos FA and Difante MVB 2013. Ingestive behavior, performance and forage intake by beef heifers on tropical pasture systems. Revista Brasileira De Zootecnia-Brazilian Journal of Animal Science 42, 549–558.

Orr DM, Burrows WH, Hendricksen RE, Clem RL, Rutherford MT, Conway MJ, Myles DJ, Back PV and Paton CJ 2001. Pasture yield and composition changes in a Central Queensland black speargrass (Heteropogon contortus) pasture in relation to grazing management options. Australian Journal of Experimental Agriculture 41, 477–485.

Orr RJ, Young KL, Cook JE and Champion RA 2005. Development of a micro-sward technique for determining intake characteristics of perennial ryegrass varieties. Euphytica 141, 65–73.

Oudshoorn FW, Cornou C, Hellwing ALF, Hansen HH, Munksgaard L, Lund P and Kristensen T 2013. Estimation of grass intake on pasture for dairy cows using tightly and loosely mounted di-and tri-axial accelerometers combined with bite count. Computers and Electronics in Agriculture 99, 227–235.

Palhano AL, Carvalho PCD, Dittrich JR, de Moraes A, da Silva SC and Monteiro ALG 2007. Forage intake characteristics on mombacagrass pastures grazed by Holstein heifers. Revista Brasileira De Zootecnia-Brazilian Journal of Animal Science 36, 1014–1021.

Penning P and Boval M 1997. Effects of fasting on ingestive behaviour of sheep grazing grass or white clover monocultures. In XVIIIth International Grassland Congress, Manitoba, Saskatchewan, Canada,

Pizzuti LAD, Alves DC, Brondani IL, Pacheco PS, Freitas LD, Segabinazzi LR, Callegaro AM and Teixeira OD 2012. Behavior pattern of beef heifers supplemented with different energy sources on oat and ryegrass pasture. Revista Brasileira De Zootecnia-Brazilian Journal of Animal Science 41, 1921–1927.

Rocha CH, Santos GT, Padilha DA, Schmitt D, Medeiros-Neto C and Sbrissia AF 2016. Displacement patterns of cattle grazing on Kikuyugrass swards under intermittent grazing. Arquivo Brasileiro De Medicina Veterinaria E Zootecnia 68, 1647–1654.

Roguet C, Prache S and Petit M 1998. Feeding station behaviour of ewes in response to forage availability and sward phenological stage. Applied Animal Behaviour Science 56, 187–201.

Rook AJ, Dumont B, Isselstein J, Osoro K, WallisDeVries MF, Parente G and Mills J 2004. Matching type of livestock to desired biodiversity outcomes in pastures - a review. Biological Conservation 119, 137–150.

Rutter SM, Orr RJ, Penning PD, Yarrow NH and Champion RA 2002. Ingestive behaviour of heifers grazing monocultures of ryegrass or white clover. Applied Animal Behaviour Science 76, 1–9.

Sichonany MJD, da Rocha MG, Potter L, da Rosa ATN, de Oliveria A, Ribeiro LA, Stivanin SCB and Alves MB 2017. Effect of supplementation frequency in feeding behavior, displacement patterns and forage intake of beef heifers. Semina-Ciencias Agrarias 38, 3215–3229.

Soca P, Gonzalez H, Manterola H, Bruni M, Mattiauda D, Chilibroste P and Gregorini P 2014. Effect of restricting time at pasture and concentrate supplementation on herbage intake, grazing behaviour and performance of lactating dairy cows. Livestock Science 170, 35–42.

Soder KJ, Gregorini P, Scaglia G and Rook AJ 2009. Dietary Selection by Domestic Grazing Ruminants in Temperate Pastures: Current State of Knowledge, Methodologies, and Future Direction. Rangeland Ecology & Management 62, 389–398.

Sprinkle JE, Holloway JW, Warrington BG, Ellis WC, Stuth JW, Forbes TDA and Greene LW 2000. Digesta kinetics, energy intake, grazing behavior, and body temperature of grazing beef cattle differing in adaptation to heat. Journal of Animal Science 78, 1608–1624.

Stakelum G and Dillon P 2004. The effect of herbage mass and allowance on herbage intake, diet composition and ingestive behaviour of dairy cows. Irish Journal of Agricultural and Food Research 43, 17–30.

Taweel HZ, Tas BM, Smit HJ, Tamminga S and Elgersma A 2006. A note on eating behaviour of dairy cows at different stocking systems-diurnal rhythm and effects of ambient temperature. Applied Animal Behaviour Science 98, 315–322.

Teixeira FA, Bonomo P, Pires AJV, da Silva FF, Marques JD and de Santana HA 2011. Displacement and permanency patterns of grazing cattle on Brachiaria decumbens deferred under four fertilization strategies. Revista Brasileira De Zootecnia-Brazilian Journal of Animal Science 40, 1489–1496.

Thanner S, Dohme-Meier F, Gors S, Metges CC, Bruckmaier RM and Schori F 2014. The energy expenditure of 2 Holstein cow strains in an organic grazing system. Journal of Dairy Science 97, 2789–2799.

Tharmaraj J, Wales WJ, Chapman DF and Egan AR 2003. Defoliation pattern, foraging behaviour and diet selection by lactating dairy cows in response to sward height and herbage allowance of a ryegrass-dominated pasture. Grass and Forage Science 58, 225–238.

Ungar ED and Griffiths WM 2002. The imprints created by cattle grazing short sequences of bites on continuous alfalfa swards. Applied Animal Behaviour Science 77, 1–12.

Ungar ED, Ravid N and Bruckental I 2001. Bite dimensions for cattle grazing herbage at low levels of depletion. Grass and Forage Science 56, 35–45.

Ungar ED, Ravid N, Zada T, Ben-Moshe E, Yonatan R, Baram H and Genizi A 2006. The implications of compound chew-bite jaw movements for bite rate in grazing cattle. Applied Animal Behaviour Science 98, 183–195.

Utsumi SA, Cangiano CA, Galli JR, McEachern MB, Demment MW and Laca EA 2009. Resource heterogeneity and foraging behaviour of cattle across spatial scales. BMC Ecology 9, 9.

Vilela HH, de Lana Sousa BM, Rozalino Santos ME, Santos AL, de Assis CZ, Rocha GdO, Faria BD and do Nascimento Junior D 2012. Forage mass and structure of piata grass deferred at different heights and variable periods. Revista Brasileira De Zootecnia-Brazilian Journal of Animal Science 41, 1625–1631.

WallisDeVries MF, Laca EA and Demment MW 1999. The importance of scale of patchiness for selectivity in grazing herbivores. Oecologia 121, 355–363.

Yayota M, Kato A, Ishida M and Ohtani S 2015. Ingestive behavior and short-term intake rate of cattle grazing on tall grasses. Livestock Science 180, 113–120.

